# 3’ Branch Ligation: A Novel Method to Ligate Non-Complementary DNA to Recessed or Internal 3’OH Ends in DNA or RNA

**DOI:** 10.1101/357863

**Authors:** Lin Wang, Yang Xi, Wenwei Zhang, Weimao Wang, Hanjie Shen, Xiaojue Wang, Xia Zhao, Andrei Alexeev, Brock A. Peters, Alayna Albert, Xu Xu, Han Ren, Ou Wang, Killeen Kirkconnell, Helena Perazich, Sonya Clark, Evan Hurowitz, Ao Chen, Xun Xu, Radoje Drmanac, Yuan Jiang

## Abstract

Nucleic acid ligases are crucial enzymes that repair breaks in DNA or RNA during synthesis, repair and recombination. Various molecular tools have been developed using the diverse activities of DNA/RNA ligases. Herein, we demonstrate a non-conventional ability of T4 DNA ligase to join 5’ phosphorylated blunt-end double-stranded DNA to DNA breaks at 3’ recessive ends, gaps, or nicks to form a 3’ branch structure. Therefore, this base pairing-independent ligation is termed 3’ branch ligation (3’BL). In an extensive study of optimal ligation conditions, similar to blunt-end ligation, the presence of 10% PEG-8000 in the ligation buffer significantly increased ligation efficiency. A low level of nucleotide preference was observed at the junction sites using different synthetic DNAs. Furthermore, we discovered that T4 DNA ligase efficiently ligated DNA to the 3’ recessed end of RNA, not to that of DNA, in a DNA/RNA hybrid, whereas RNA ligases are less efficient in this reaction. These novel properties of T4 DNA ligase can be utilized as a broad molecular technique in many important applications. We performed a proof-of-concept study of a new directional tagmentation protocol for next generation sequencing (NGS) library construction that eliminates inverted adapters and allows sample barcode insertion adjacent to genomic DNA. 3’BL after single transposon tagmentation can theoretically achieve 100% usable template, and our empirical data demonstrate that the new approach produced higher yield compared with traditional double transposon or Y transposon tagmentation. We further explore the potential use of 3’BL for preparing targeted RNA NGS libraries with mitigated structure-based bias and adapter dimer problems.

## Introduction

Ligases repair breaks in nucleic acids, and this activity is essential for cell viability and vitality. DNA ligases catalyze the formation of a phosphodiester bond between DNA ends and play crucial roles in DNA repair, recombination, and replication *in vivo*^1–3^. RNA ligases join the 5’-phosphoryl (5’PO_4_) and 3’-hydroxyl (3’OH) RNA termini via phosphodiester bonds and are involved in RNA repair, splicing, and editing^4^. Ligases from all three kingdoms of organisms (bacteria, archaebacteria, and eukaryotes) can be utilized *in vitro* as important molecular tools for applications such as cloning, ligase-based amplification or detection, and synthetic biology^5–7^. One of the most widely used ligases *in vitro* is bacteriophage T4 DNA ligase, which is a single 55-kDA polypeptide that requires ATP as an energy source^8^. T4 DNA ligase typically joins the adjacent 5’PO_4_ and 3’OH termini of duplexed DNA. In addition to sealing nicks and ligating cohesive ends, T4 DNA ligase can also efficiently catalyze blunt-end joining, which has not been observed for any other DNA ligases^9,10^. Some unusual catalytic properties of this ligase were reported previously, such as sealing single-stranded gaps in duplex DNA, sealing nicks adjacent to abasic sites in double-stranded DNA (dsDNA), promoting intramolecular loop formation with partially double-stranded DNA, and joining DNA strands containing 3’ branch extensions^11–13^. Researchers also observed template-independent ligation mediated by T4 ligase, such as mis-paired nick sealing in dsDNA^14^ or even single-stranded DNA (ssDNA) ligation, albeit at very low efficiency^15^. These results suggest that perfect complementary base pairing at or adjacent to the ligation junction is not critically needed for some unconventional T4 DNA ligase activities. T4 RNA ligases 1 and 2 are the products of genes 63 and 24, respectively, of T4 phage. Both require an adjacent 5’PO_4_ and 3’OH end for a successful ligation with the concurrent hydrolysis of ATP to AMP and PPi. The substrates for T4 RNA ligase 1 include single-stranded RNA and DNA, whereas T4 RNA ligase 2 preferentially seals nicks on dsRNA rather than ligating the ends of ssRNA^16,17^.

Here, we demonstrate a non-conventional end-joining event mediated by T4 DNA ligase that we call 3’-branch ligation (3’BL). This method can join DNA or DNA/RNA fragments at nicks, single-stranded gaps, or 3’ recessive ends to form a branch structure. This report includes extensive study of a wide variety of ligation cofactors and activators and the optimization of the ligation conditions for this type of novel ligation. With our 3’BL protocol, no base pairing was required, and the ligation can reach 70-90% completion in most cases, including for a 1-nt gap. One application of this method is the attachment of adapters to DNA or RNA during NGS library preparation. Several genomic structures that were previously considered unligatable can now be used as substrates for 3’BL, resulting in a high conversion rate of input DNA into adapter-ligated molecules while avoiding chimeras. We demonstrate that 3’BL can be coupled with transposon tagmentation to increase library yield. The directional tagmentation strategy we propose will theoretically produce templates 100% of which can be utilized for sequencing. Our study demonstrated the value of this novel technique for NGS library preparation and the potential to advance many other molecular applications.

## Results

### 3’ Branch ligation: a novel method to ligate non-complementary DNA ends

Conventionally, DNA ligation involves the joining of the 5’PO_4_ and 3’OH DNA ends of cohesive or blunt-ended fragments. Cohesive-end ligation is generally faster and less dependent on enzyme concentration compared with blunt-end joining. Both processes can be catalyzed by bacteriophage T4 DNA ligase, which uses ATP as an energy-yielding cofactor and requires Mg^2+8^. T4 DNA ligase was also reported to ligate specific or degenerate single-stranded oligos to partially single-stranded substrates through hybridization^18,19^. Here, we demonstrated a non-conventional T4 DNA ligase-mediated ligation that does not require complimentary base pairing and can ligate a blunt-end DNA donor to the 3’OH end of a duplex DNA acceptor at 3’ recessed strands, gaps, or nicks (Figure 1a). Therefore, we use the term 3’-branch ligation (3’BL) to describe these ligations. The synthetic donor DNA we used contained a 5’ blunt duplex end and a 3’ ssDNA end. The acceptor substrates contained one of the following structures: a dephosphorylated nick, a 1- or 8-nucleotide (nt) gap, or a 3’ 36-nt recessed end (Supplementary Table 1). T4 ligase helps to join the 5’PO4 of the donor strand to the sole ligatable 3’OH of the acceptor strand to form a branch-shaped ligation product.

**Figure 1.**
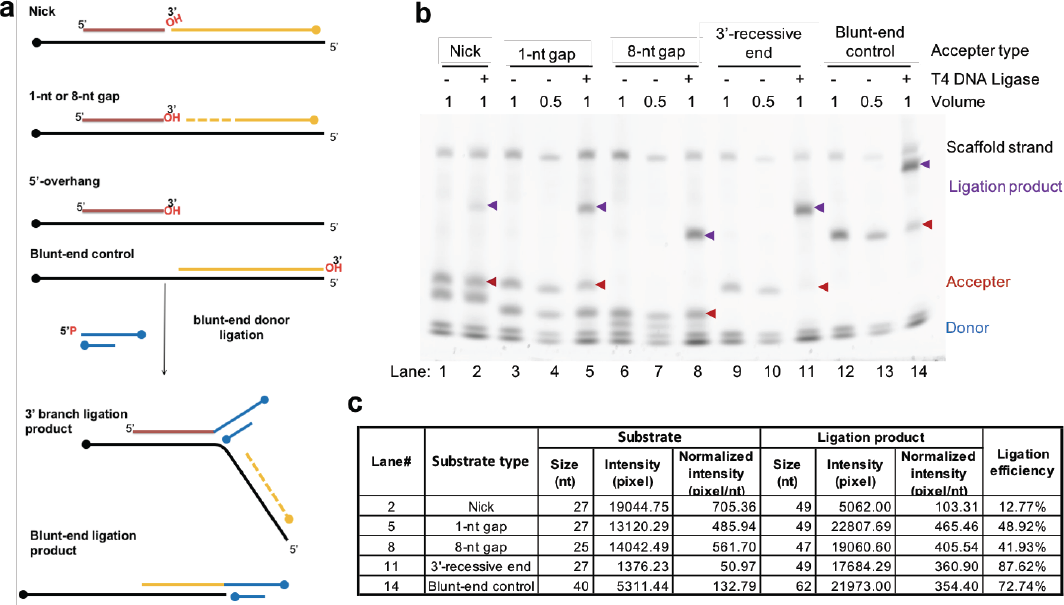
3’ branch ligation by T4 DNA ligase at non-conventional DNA ends formed by nicks, gaps, and overhangs. (a) Schematic representation of ligation assay with different DNA accepter types. The blunt-end DNA donor (blue) is a synthetic, partially dsDNA molecule with dideoxy 3’-termini (filled circles) to prevent DNA donor self-ligation. The long arm of the donor is 5’-phosporylated. The DNA acceptors were assembled using 2 or 3 oligos (black, red, and orange lines) to form a nick (without phosphates), a gap (1 or 8 nt), or a 36-nt 3’-recessive end. All strands of the substrates are unphosphorylated, and the scaffold strand is 3’ dideoxy protected. (b) Analysis of the size shift of ligated products of substrates 1, 2, 3, and 4, respectively, using a 6% denaturing polyacrylamide gel. The negative no-ligase controls (lanes 1, 3, 4, 6, 7, 9, 10, 12, and 13) were loaded at 1 or 0.5X volume of corresponding experimental assays. If ligation occurs, the substrate size is shifted up by 22 nt. Red arrowheads correspond to the substrate, and purple arrowheads correspond to donor-ligated substrates. Thermo Fisher’s 25-bp DNA Ladder was used. Donor and substrate sequences in Table S1. (c) Expected sizes of substrate and ligation product and approximate ligation efficiency in each experimental group. The intensity of each band was estimated using ImageJ and normalized by its expected size. Ligation efficiency was estimated by dividing the normalized intensity of ligated products by the normalized total intensity of ligated and unligated products.

To optimize the ligation efficiency, we extensively tested a number of factors that affect general ligation efficiency, including the adapter:DNA substrate ratio, T4 ligase quantity, final ATP concentration, Mg^2+^ concentration, pH, incubation time, and different additives, such as polyethyleneglycol-8000 (PEG-8000) and single-stranded binding protein (SSB) (Supplementary Figure 1 and 2, data not shown). Adding PEG-8000 to a final concentration of 10% substantially increased the ligation efficiency from less than 10% to more than 80% (data not shown, Figure 1 and 2). A large range of ATP concentrations (from 1 µM to 1 mM in Figure S1a) and Mg^2+^concentrations (3 mM to 10 mM, data not shown) were compatible with the 3’BL. The ligase quantity needed for 3’BL was comparable to that for blunt-end ligation. In our optimized conditions, we used donor:substrate DNA molar ratios of 30 to 100, and we performed the reactions at 37°C for one hour at pH 7.8 with 1 mM ATP, 10 mM MgCl_2_, and 10% PEG-8000. The ligation of the same donors to blunt-end substrates and ligase-free reactions were used as positive and negative controls, respectively. The ligation donor (Ad-G) is double-stranded on one end (5’ phosphorylated and 3’ dideoxy protected) and single-stranded (3’ dideoxy protected) on the other end (Fig. 1a, Table S1). The ligation substrates are composed of the same bottom strand (ON1) with different top strands to compose nick, gap, and overhang structures. To quantify ligation product yields, the reaction products were separated on 6% denaturing polyacrylamide gels (Figure 1b). Ligation efficiency was calculated as the ratio of product to substrate intensity using ImageJ (Figure 1b-c). The 3’-recessive ligation (lane 11 in Figure 1b) appeared approximately 90% complete, which is even higher than the blunt-end ligation control (lane 14, 72.74%) and suggests remarkably high ligation efficiency with 3’-recessive DNA ends. The 1- or 8-nt gap substrates (lane 5 and 8) showed good ligation efficiency of approximately 45%. Nick ligation (lane 2) efficiency was the lowest at approximately 13%. However, this ligation yield was improved when the nick ligation reaction was incubated longer (data not shown), suggesting slower kinetics for the nick ligation reaction.

We also extended our study to different adapter and substrate sequences (Figure 2). The 5’PO_4_ends of three different adapters (Ad-T, Ad-A, or Ad-GA in Supplementary Table 1) contained either a single T or A or the dinucleotide GA at the ligation junction before a consensus CTGCTGA sequence. These 5’PO_4_ ends were individually ligated to the 3’OH ends of acceptor templates with a T at the ligation junction. Overall, high ligation efficiency (70-90%) was seen in most cases except for nick ligations or 3’BL using Ad-GA (Figure 2f), thus indicating some nucleotide preference of T4 DNA ligase at the ligation junctions. Independent of the adapter and substrate sequences, the 3’-recessive end or gap ligations always showed better efficiencies (60-90%), whereas the nick ligation was fairly inefficient, in a one-hour incubation. We hypothesize that these discrepancies in ligation efficiency are due to the DNA bending where the nick/gap/overhang starts and exposes the 3’OH group for ligation. The longer ssDNA region likely makes the 3’ termini more accessible in the ligation and results in higher ligation efficiency. We also tested whether a similar end-joining event could occur as 5’ branch ligation. In contrast to 3’BL, no obvious ligation of a blunt-end adapter to the 5’PO_4_ end at the gap or the 5’-recessive end was observed (data not shown). This result suggests greater steric hindrance of T4 DNA ligase at the donor’s 5’ termini compared with 3’ termini.

**Figure 2.**
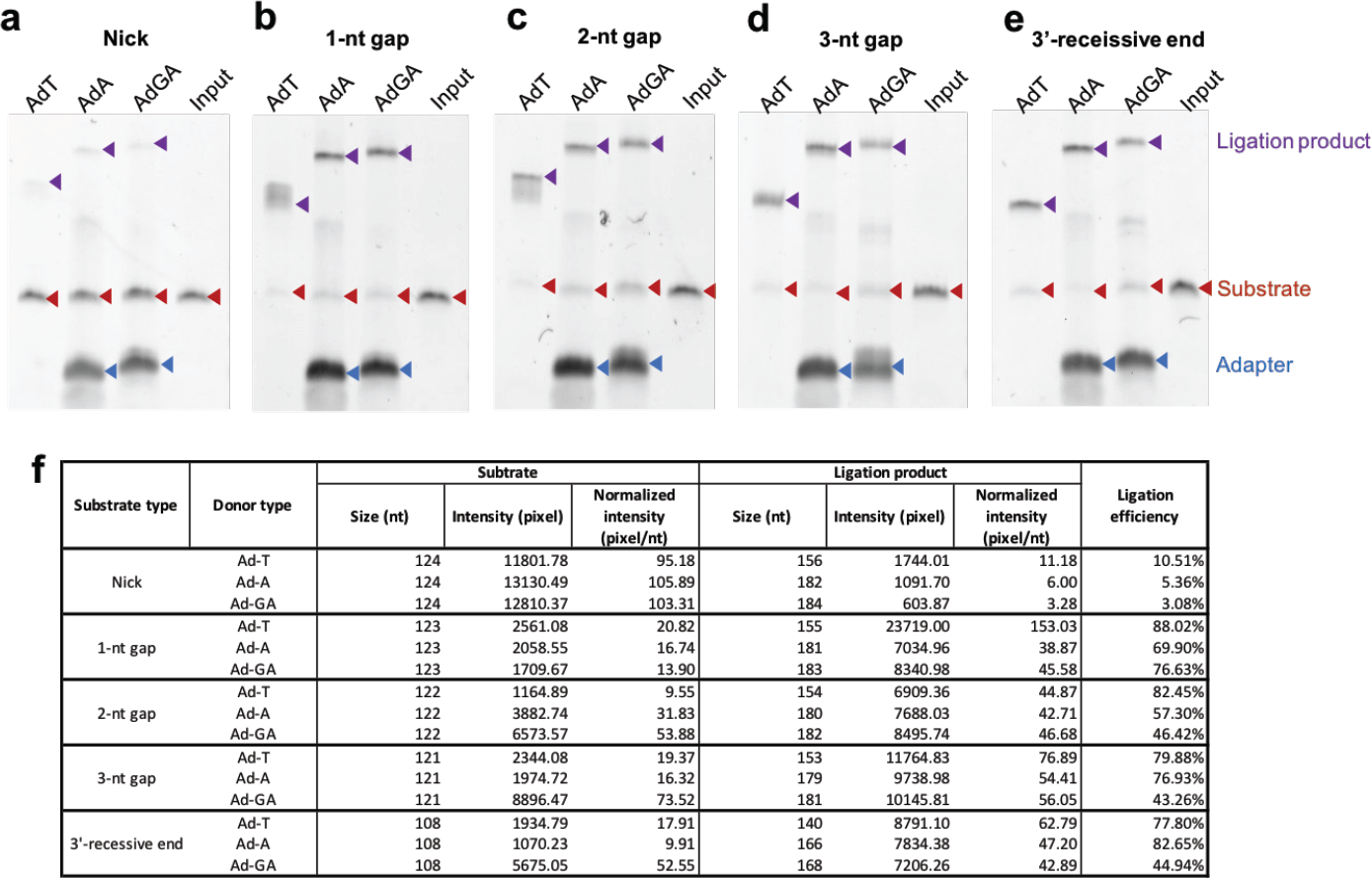
Gel analysis of size shift of ligated products using 6% TBE polyacrylamide gel. Red arrowheads correspond to the substrate, and purple arrowheads correspond to donor-ligated substrates: substrate 5 (nick) (a), substrate 6 (1-nt gap) (b), substrate 7 (2-nt gap) (c), substrate 8 (3-nt gap), (d) and substrate 9 (3’-recessive end) (e). Thermo Fisher’s 25-bp DNA Ladder was used. Three DNA donors with different bases at the 5’-end of the ligation junction (T, A, or GA) were examined. Donor and substrate sequences are summarized in Table S2. (f) Table of ligation efficiency calculated based on normalized band intensity using ImageJ.

### 3’ Branch ligation to ligate DNA to RNA

We further investigated 3’BL on DNA/RNA hybrids (ON-21/ON-23 in Table S1) that form one DNA and one RNA 5’-overhang (Figure 3a). Ligation on DNA/DNA hybrids served as a positive control, whereas negative ligation controls included DNA/RNA hybrids, ssDNA, or ssRNA oligos incubated individually or with adapters (lanes 3, 4, and 5 in Figure 3c and data not shown). Interestingly, when DNA/RNA hybrids were incubated with a blunt-end dsDNA donor, we observed a size change of the RNA oligo from the original 29 nt to 49 nt upon ligation. However, the DNA substrate remained unchanged (lanes 1 and 2 in Figure 3c). This result suggests that the blunt-end dsDNA donor ligated to the 3’-end of RNA at the 3’-recessive DNA end but not to the 3’-end of DNA at the 3’-recessive RNA termini. As a positive control, DNA/DNA hybrids with 3’ recessive ends on each side showed band shifts to larger species on both strands with nearly 100% efficiency (data not shown). To confirm that the 3’ recessive structure was needed for 3’BL, we performed the same ligation reaction while replacing the original DNA oligo (ON-21) with another long DNA template (ON-23) that is not complimentary to ON-22 RNA (Figure 3b). Unsurprisingly, no ligation was observed using the ON-23 DNA template (lane 10-13 in Figure 3c). Our finding indicates that T4 DNA ligase can promote 3’BL on DNA/RNA hybrids and that this activity has certain steric substrate preferences that may be affected by differences in T4 DNA ligase-substrate binding affinities.

**Figure 3:**
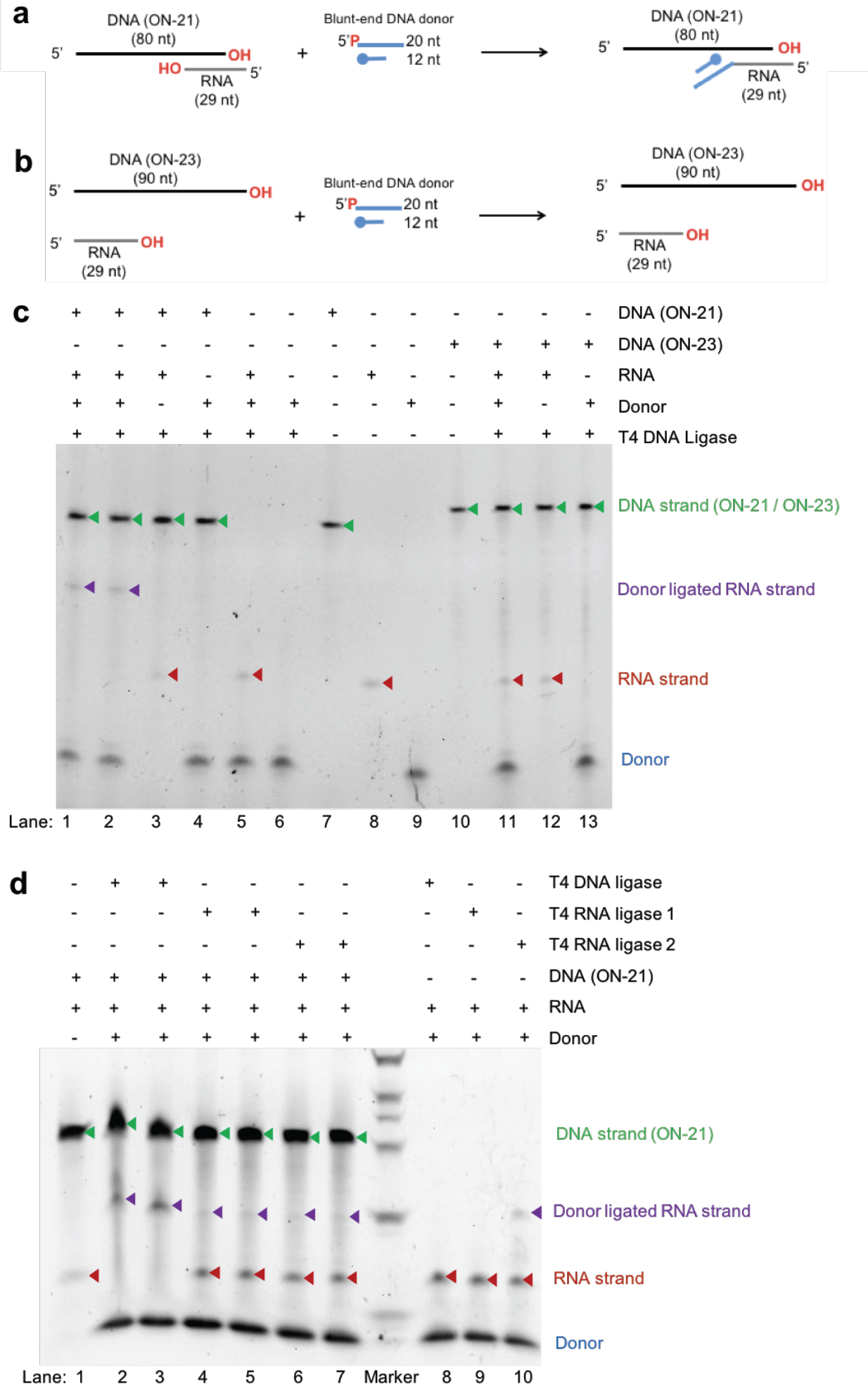
3’ branch ligation at the 3’ end of RNA in DNA/RNA hybrid. Schematic representation of 3’-branch ligation on a DNA/RNA hybrid with a 20-bp complimentary region. We tested whether blunt-end DNA donors would ligate to the 3’-recessive end of DNA and/or to the 3’-recessive end of RNA. DNA(ON-21) hybridizes with the RNA strand (a), whereas DNA(ON-23) cannot hybridize with the RNA strand (b). (c, d) Gel analysis of size shift of ligated products using 6% denaturing polyacrylamide gel. The red arrowheads correspond to the RNA substrate (29 nt), and the green arrowhead corresponds to DNA substrate (80 nt). The purple arrowhead corresponds to donor-ligated RNA substrates. If ligation occurs, the substrate size would shift up by 20 nt. (c) Lane 1 and 2, experimental duplicates; lanes 7-10, no-ligase controls; 10% PEG was added with T4 DNA ligase. (d) Lane 1, no-ligase control; lane 2, 3, and 8, T4 DNA ligase with 10% PEG; lane 4, 5, and 9, T4 RNA ligase 1 with 20% DMSO; lane 6, 7, and 10, T4 RNA ligase 2 with 20% DMSO. Thermo Fisher’s 25-bp DNA Ladder was used.

A previous study reported that for sealing nicks in DNA/RNA hybrids, T4 DNA ligase and T4 RNA ligase 2, but not T4 RNA ligase 1, can effectively join a 5’PO_4_ DNA end to a juxtaposed 3’OH DNA or RNA end when the complimentary strand is RNA but not DNA^17^. Therefore, we performed the same ligation test using T4 RNA ligase 1 and 2 either in 20% DMSO (Figure 3d) or in 10% PEG (data not shown). In both tests, T4 RNA ligase 1 and T4 RNA ligase 2 slightly ligated the blunt end adapters to the 3’ end of RNA in a DNA/RNA hybrid. Notably, in the RNA-only controls, T4 RNA ligase 2 could join a blunt-end dsDNA adapter to ssRNA. In conclusion, T4 DNA ligase, but not T4 RNA ligase, is competent to efficiently ligate blunt-end dsDNA to the 3’ end of RNA through 3’BL.

### Directional tagmentation library construction

Because 3’BL is useful for ligating adapters to several genomic structures with high efficiency, we explored its application in NGS workflows. Transposon-based library construction is rapid and consumes less input DNA compared with conventional NGS library preparation. However, using commercial transposon-based library preparation systems, only half of tagged molecules are flanked by two different adapter sequences (Figure 4a), and tagged DNA is flanked by self-complementary regions that could form stable hairpin structures and compromise sequencing quality^20^. In addition, the PCR-mediated incorporation of adapter sequences has not been adapted for whole-genome bisulfite sequencing or PCR-free NGS library construction.

To overcome these limitations, we developed a new protocol for transposon-based NGS library construction by incorporating 3’BL. Both Tn5 and MuA transposons work through a “cut and paste” mechanism, in which a transposon adapter sequence is end-joined to the 5’-end of target DNA to create a 9-bp or 5-bp gap, respectively, at the 3’-end of the genomic DNA (Figure 4a). Subsequently, 3’BL can be used to add another adapter sequence to the 3’-end of genomic DNA at the gap to complete the directional adapter ligation (Figure 4c). We used Tn5 transposons in this manuscript to compare the efficiency of the single-tagmentation + 3’BL approach (Figure 4c) to that of a double-tagmentation approach, which uses the two different Tn5-based adapters TnA and TnB (Figure 4a), and to that of another directional single-tagmentation strategy using Y adapters that contain two different adapter sequences (Figure 4b). Human genomic DNA was incubated with equimolar amounts of TnA and TnB transposome complexes, with the TnA transposome complex alone, or with the TnY (TnA/B) transposome complex. The product of TnA transposome-only fragmentation was further used as a template for 3’BL with the blunt-end adapter AdB, which shares a common adapter sequence with TnB. PCR amplification was performed using two primers, Pr-A and Pr-B, designed to recognize the TnA and AdB/TnB adapters, respectively. The quantification data suggested that TnA&AdB had the highest efficiency compared to TnA&TnB and TnY (TnA/B) (Figure 4d). No significant amplification was observed when only one primer specific to TnA adapter was used (Figure 4d). As expected due to PCR suppression, the TnA-3’BL approach, the double-tagmentation approach, and the TnY approach all showed significantly higher PCR efficiency than the tagmentation reaction with only the TnA or TnB transposome complex alone (Figure 4d).

**Figure 4.**
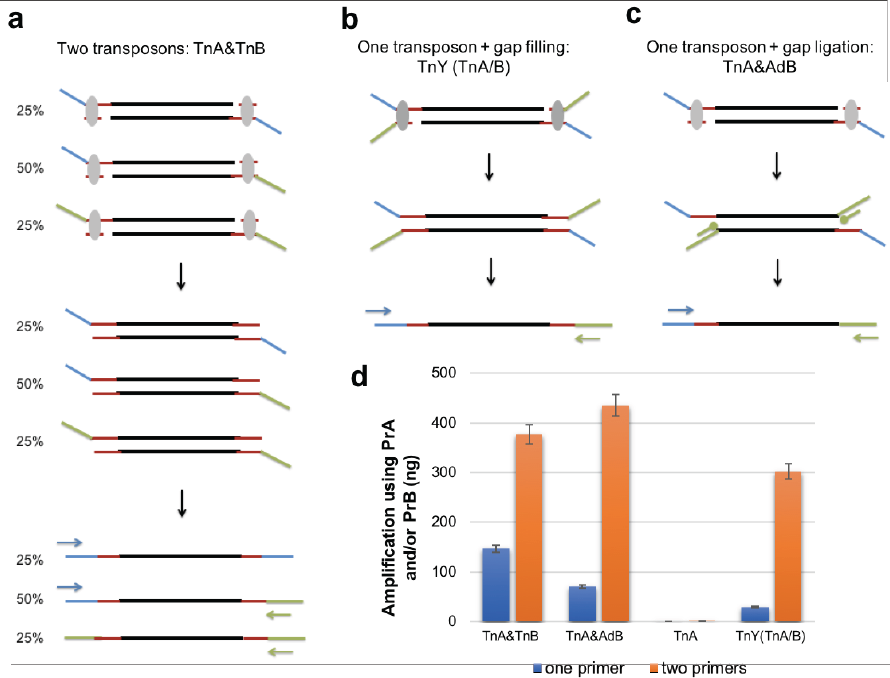
(a-c) Schematic representation of three transposon tagmentation methods followed by PCR amplification using Pr-A (blue arrow) and Pr-B (green arrow). Two-transposons method (a); one Y transposon tagmentation with 3’-gap filling (b); One-transposon method with adapter ligation at 3’-gap (c). (d) Graph of amplification signal after purification using pr-A or pr-A with pr-B after the various tagmentation and gap ligation conditions.

We also sequenced these libraries using BGI’s DNA nanoball bases sequencing platform and compared the base-positional bias among the transposon-interfered end, the 3’BL end, and the regular TA ligation end (Figure 5). It is obvious that the positional bias at the 3’BL end is less than that at the Tn5 end (Figure 5a-b), which occurs because the 3’BL end is influenced by both transposon interruption and 3’BL. Because only the first 6 nt (position 1-6) of the 3’BL end showed base bias and the bias was similar to but not exactly the same as that of its hybridized Tn5 end (position 30-35, after the 9-nt overhang), we conclude that the positional bias we observed at the 3’BL end is mainly caused by the Tn5 transposon. Therefore, 3’BL causes minimal bias and is similar to regular TA ligation (Figure 5c).

**Figure 5.**
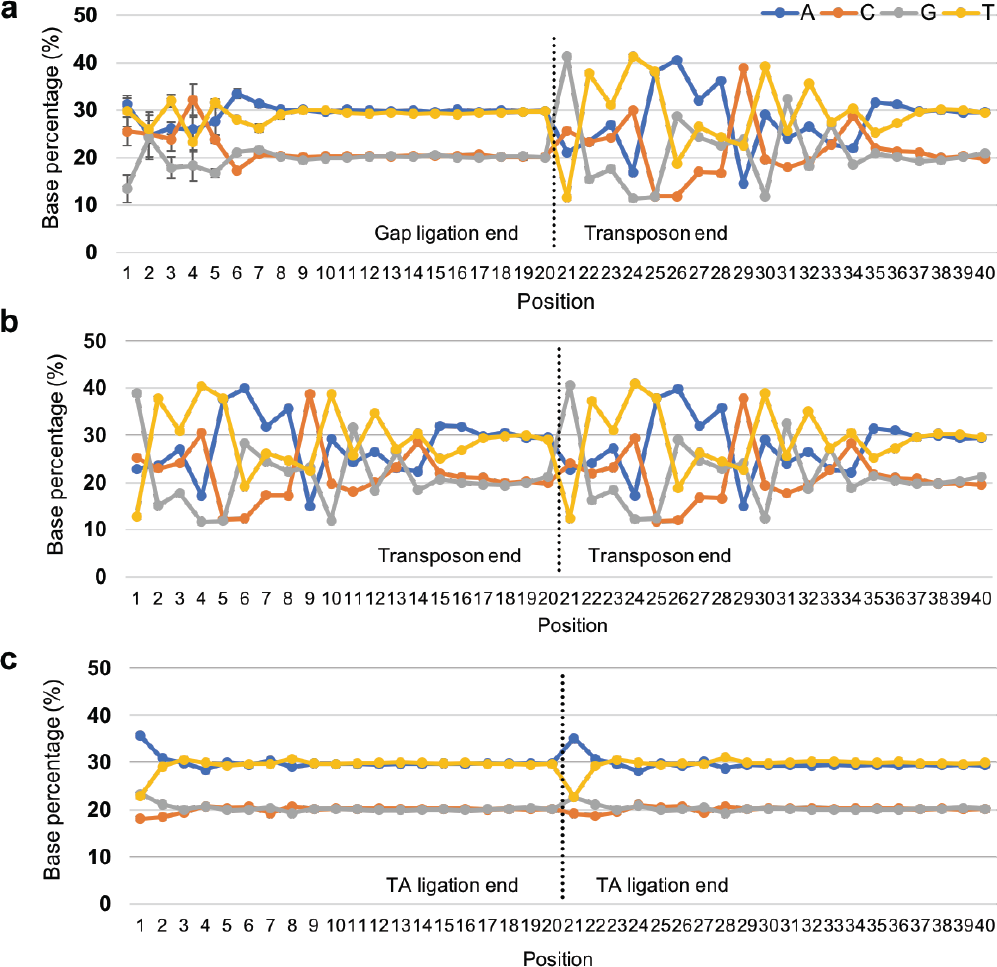
Base distribution bias of Tn5-gap ligation (a), two transposons (b), and regular TA ligation (c). Only the first 20 bases from each end of the ligation are presented; adenine, blue; cytosine, orange; guanine, gray; thymine, yellow; the average and standard deviation of five independent libraries are presented.

## Discussion

One important property of T4 DNA ligase is its efficient joining of blunt-ended dsDNA^21,22^, which has not been observed with other DNA ligases. This ligase was also reported to mediate some unusual catalytic events, such as ligating single-strand gaps or mismatched bases in duplex DNA^11,12^, forming a stem-loop molecule from partially double-stranded DNA^13^, or inefficiently ligating ssDNA in a template-independent manner^15^.

Here, we demonstrated that T4 DNA ligase catalyzed the joining of blunt-end dsDNA to the 3’OH end of dsDNA with a nick and the joining of partially single-stranded duplex DNA with a gap or 5’ overhang. In contrast, no ligation to the 5’PO4 end at the 5’ recessed ends or in the gaps was observed (data not shown). This indicates that after binding to the 5’PO4 end of the dsDNA adapter, T4 DNA ligase can access the recessed 3’ end in the target DNA when DNA bends and exposes the 3’OH end. Unlike the mismatched or gapped ligation in previous reports^11–14^, no base pairing of the donor DNA to the acceptor DNA was required with our 3’BL method. Moreover, even for a 1-nt gap, greater than 70% completion was accomplished using optimized conditions, while other template-independent ligation was of very low yield and was only detected by qPCR measurement^15^.

However, different 3’BL ligation efficiencies were observed for ligating 5’ T, A, or GA to 3’ T (Figure 2), which indicates some sequence preference at the ligation junction. Despite recognized ligation bias^23^, T4 DNA ligase is commonly used in the adapter addition step during NGS library preparation. With its ability to perform 3’BL, T4 ligase can ligate adapters to several genomic structures that were previously considered unligatable, resulting in a higher template usage rate. For example, 3’BL can be coupled with single transposon tagmentation.

The traditional double-transposon strategy has only 50% of the tagmented molecules that are amenable for the following amplification step. However, when DNA tagmentation is performed using one transposon with subsequent 3’BL, an increased yield of molecules with different adapters on each insert end can be acquired (Figure 4). Furthermore, the tagmented-3’BL products can be directly loaded on Illumina’s flow cell as PCR free WGS libraries, which was difficult to achieve using double-transposon strategy. Other directional transposon protocols have been proposed using a Y transposon composed of two different adapter sequences^24^ or replacing the unlinked strand from a single transposon with a second adapter oligo followed by gap filling and ligation^25^. We also tested ligation of a single-stranded adapter oligo with 6-8 degenerated bases at 5’ end and obtained similar yield as with 3’BL (data not shown). However, these approaches continue to preserve the inverted adapter sequences and/or cannot insert sample barcodes adjacent to genomic DNA as the tagmented-3’BL protocol can. Based on NGS data, the 3’BL ligated genomic ends also demonstrated fewer positions with positional base composition bias, and the first 6-nt bias was mild and mainly caused by transposon insertion, suggesting that 3’BL has minimal positional bias. Using this new library construction method, Wang et al. successfully achieved highly accurate and complete variant calling in WGS and near-perfect phasing of variants into long contigs with N50 size up to 23.4 Mb for long fragment reading (BioRxiv, https://doi.org/10.1101/324392).

In this study, we also investigated 3’BL using templates of a chimeric DNA/RNA duplex that forms a 5’ DNA and a 5’ RNA overhang (Figure 3). Unexpectedly, blunt-ended dsDNA was efficiently ligated to the 3’ termini of RNA, but not DNA, suggesting that T4 ligase has a ternary complex formation preference. The ligation efficiency was greatly reduced if T4 RNA ligase I or II was used to join the ends. Another preliminary yet important application of 3’BL with T4 DNA ligase is the enrichment of mRNA or the construction of targeted RNA libraries, especially for miRNAs, the small regulatory RNAs whose uncontrolled expression leads to a number of diseases^26,27^. Thus, our 3’BL technique can be readily applied to the detection of cancer and Alzheimer’s disease using miRNA. Hybridization with DNA probes targeting the Poly(A) tail or specific miRNA sequences can be used to create DNA-RNA hybrids with a DNA 5’-overhang, which is followed by ligation to adapter sequences with sample and/or UID barcodes through 3’BL. These common sequences can then be reverse transcribed to produce the cDNA of targeted RNA sequences. Compared with current miRNA capture technologies, the use of 3’BL mediated by T4 DNA ligase could potentially provide several advantages for NGS RNA library construction. First, hybridization with a DNA strand would prevent secondary structure formation by the RNA strand and therefore mitigate the bias introduced by other protocols. Second, T4 DNA ligase enables high-efficiency adapter addition through 3’BL, which avoids intramolecular RNA interactions that can be promoted by RNA ligases. Third, adapter dimers can be effectively eliminated, possibly rendering undesirable gel purification unnecessary. This new method could lead to improved unbiased microRNA expression profiling with simple and scalable workflows, and thus, large-scale research studies would become more affordable.

The findings of this study add to the growing understanding of T4 DNA ligase activities. We envision 3’ branch ligation becoming a general tool in molecular biology that will advance the development of new DNA engineering methods beyond described NGS applications.

## Materials and Methods

### 3’-branch ligation for duplex DNA

The substrates for 3’BL were composed of 2 pmol of ON1 or ON9 mixed with 4 pmol each of one or two additional oligos in pH 8 Tris-EDTA (TE) buffer (Life Technologies) as follows: substrate 1 and 5 (nick), ON-1/2/3 and ON-9/10/11; substrate 2 and 6 (1-nt gap), ON1/2/4 and ON9/10/12; substrate 3 (8-nt gap), ON1/4/5; substrate 4 and 9 (5’ overhang), ON1/2 and ON9/10; substrate 7 (2-nt gap), ON9/10/13; substrate 8 (3-nt gap), ON9/10/14; blunt-end control, ON1 and ON6 (Figure 1, Supplementary Table 1). The template was ligated to 180 pmol of adapter (Ad-G: ON7/8, Ad-T: ON15/16, Ad-A: ON17/8, or Ad-GA: ON19/20) using 2,400 units of T4 ligase (Enzymatics Inc.) in 3’BL buffer [0.05 mg/ml BSA (New England Biolabs), 50 mM Tris-Cl pH 7.8 (Amresco), 10 mM MgCl_2_ (EMD Millipore), 0.5 mM DTT (VWR Scientific), 10% PEG-8000 (Sigma Aldrich), and 1 mM ATP (Sigma Aldrich)]. The optimization tests were conducted by altering the ATP concentration from 1 µM to 1 mM, the Mg^2+^ concentration from 3 to 10 mM, the pH value from 3 to 9, temperature from 12 to 42°C, and adjusting additives such as PEG-8000 from 2.5% to 10% and SSB from 2.5 to 20 ng/µL. The ligation mixture was prepared on ice and incubated at 37°C for 1 to 12 hours before heat inactivation at 65°C for 15 min. The samples were purified using Axygen beads (Corning) and eluted into 40 µL TE Buffer. All ligation reactions were run on 6% TBE or denaturing polyacrylamide gels (Life Technologies) and visualized on an Alpha Imager (Alpha Innotech). An input control was loaded at either an equal quantity or half the quantity of the template used for ligation. A ligation efficiency rate was estimated by dividing the intensity of ligated products by the total intensity of ligated and unligated products using ImageJ Software (NIH).

### 3’-branch ligation for DNA/RNA hybrid

The substrates for 3’BL were composed of 10 pmol ON-21 RNA oligo mixed with 2 pmol of ON-21 or ON-23 DNA oligo. For T4 DNA ligase-mediated 3’BL, the substrate was incubated with Ad-T (ON15/16) in 3’BL buffer as described above and incubated at 37°C for 1 hour. 3’BL using T4 RNA ligase 1 or 2 was performed in 1x RNA ligase buffer (NEB) with either 20% DMSO or 25% PEG. All ligation products were assayed on 6% denaturing polyacrylamide gels.

### Directional tagmentation library construction

The transposon oligonucleotides used in this experiment were synthesized by Sangon Biotech. For the 2 transposon experiments using TnA/TnB, oligos for TnA (ON24), TnB (ON25), and MErev (ON26) were annealed at a 1:1:2 ratio. For the single transposon experiment with TnA, ON24 and ON26 were annealed at a 1:1 ratio. For the Y (TnA&TnB) transposon experiment, ON24 and ON27 were annealed at a 1:1 ratio.

Transposon assembly was performed by mixing 100 pmol of pre-annealed adapters, 7 µL of Tn5 transposase, and sufficient glycerol to obtain a total 20-µL reaction, which was incubated at 30°C for 1 hour. Tagmentation of genomic DNA (Coriell 12878) was performed in 20-µL reactions containing 100 ng of gDNA, TAG buffer (homemade), and 1 µL of the assembled transposon. The reaction was incubated at 55°C for 10 min; 40 µL of 6 M guanidine hydrochloride (Sigma) was then added to remove the transposon complex from tagmented DNA, and DNA was purified using Agencourt AMPure XP beads (Beckman Coulter). The gap ligation of AdB (ON28 and ON29) to the tagmented DNA was performed at 25°C for 1 hour in reactions containing 100 pmol of the adapter, 600 U of T4 DNA ligase (Enzymatics Inc.), and 3’BL buffer. Reactions were purified using AMPure XP beads. PCR amplification of tagmented and gap-ligated DNA was performed in 50-µL reactions containing 2 µL of the tagmented or gap-ligated DNA, TAB buffer, 1 µL TruePrep Amplify Enzyme (Vazyme), 200 mM dNTPs (Enzymatics Inc.), and 400 mM each of primers Pr-A and Pr-B. Tagmented reactions were incubated as follows: 72°C for 3 min; 98°C for 30 sec; 8 cycles of 98°C for 10 sec, 58°C for 30 sec, and 72°C for 2 min; and 72°C for a 10-minute extension. Gap-ligated reactions were run using the same program without the initial 3 min extension at 72°C. PCR reactions using either prA (ON30) or both prA and prB (ON31) were purified using AMPure XP beads. Purified products were quantified using the Qubit High-Sensitivity DNA kit (Invitrogen).

## Acknowledgments

We would like to acknowledge the ongoing contributions and support of all Complete Genomics and BGI-Shenzhen employees, especially Benjamin Allred for commenting the manuscript. This work was supported in part by the Shenzhen Peacock Plan (No. KQTD20150330171505310). Employees of BGI and Complete Genomics have stock holdings in BGI.

---

**Figure S1:**
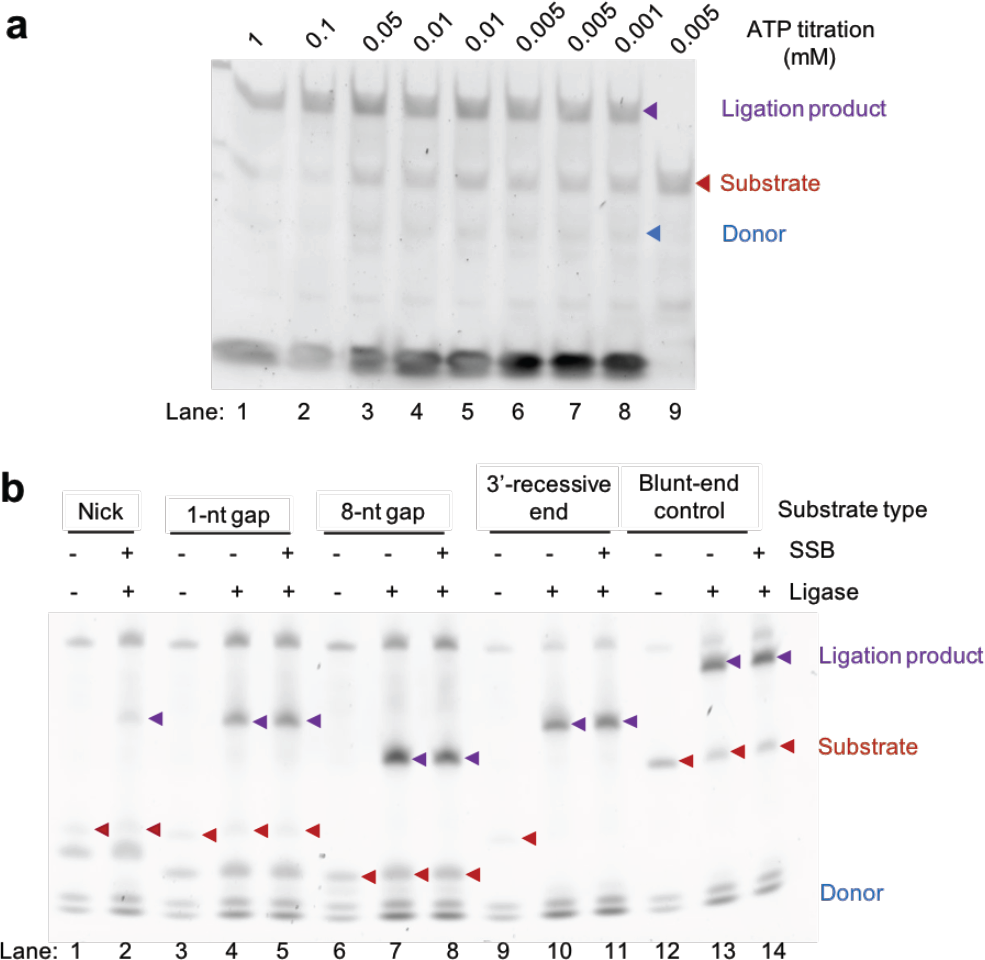
DNA 3’ branch ligation with different additive conditions. (a) Ligation at 5’-overhang DNA at titrated ATP concentrations. Duplicates were performed for 0.01 mM (lane 4 and 5) and 0.005 mM ATP (lane 6 and 7). Lane 9 is a no-donor control. (b) 3’ branch ligation of DNA at nick, 1-nt gap, 8-nt gap, 5’-overhang, and blunt end with or without SSB and ligase. Red arrowheads correspond to the substrate, and purple arrowheads correspond to donor-ligated substrates.

**Table S1.**
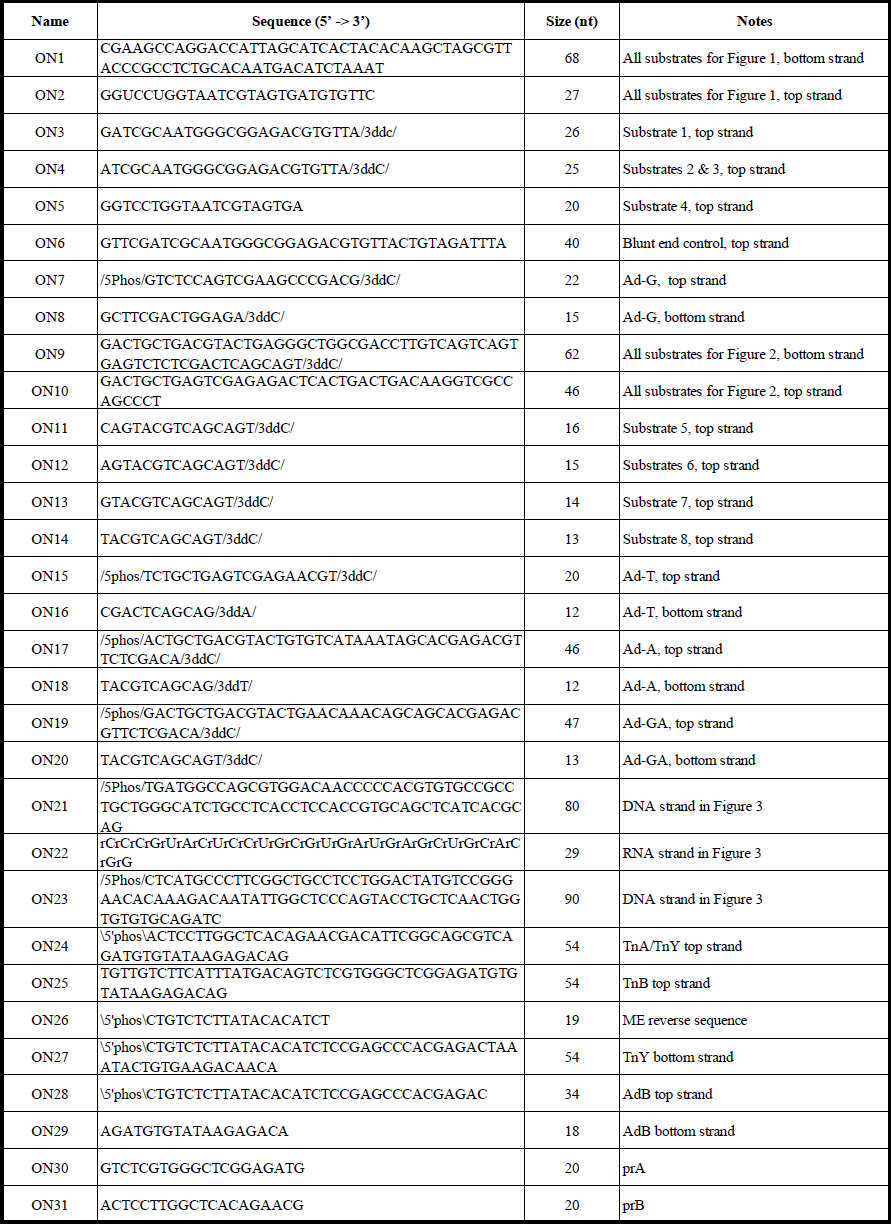
DNA donor and substrate sequences used in this manuscript.

